## Supplementary Material S1 for "Contributions of mechanical loading and hormonal changes to eccentric hypertrophy during volume overload: a Bayesian analysis using logic-based network models"

(3) Birmingham Veterans Affairs Health Care System, Birmingham, Alabama, United States of America.

(4) Division of Cardiovascular Disease, Heersink School of Medicine, University of Alabama at Birmingham, Birmingham, Alabama, United States of America.

(5) Division of Cardiothoracic Surgery, Heersink School of Medicine, University of Alabama at Birmingham, Birmingham, Alabama, United States of America.

### S1.1. Sources of experimental data from canine MVR and experimental VO in rats used to fit the network model of cardiomyocyte hyperthrophy.

**Table S1.1.** Data sources for untreated VO.

| Experimental measurement | Sources | |
|---|---|---|
| | Canine MVR | VO in rats |
| **Growth at cell and organ level** | | |
| **LV mass/BW** | (Carabello et al., 1992; Dell'Italia et al., 1995, 1997; Hankes et al., 2006; Katayama et al., 1988; Kihara et al., 1988; Matsuo et al., 1998; Nakano et al., 1991; Nemoto et al., 2002, 2005; Pat et al., 2008, 2010; Perry et al., 2002; Sabri et al., 2008; Tallaj et al., 2003; Tsutsui et al., 1994) | (Bauer et al., 1997; Gardner et al., 2010; Isgaard et al., 1994; Ishiye et al., 1995; Jobe et al., 2009; Kristen et al., 2006; Lachance et al., 2014; Lu et al., 2012; Mclarty et al., 2012; Oka et al., 1993; Pu et al., 2009; Ruzicka et al., 1993; Shuji et al., 2000; Stefano et al., 2006; Wang et al., 2012; Willenbrock et al., 1997; Zhang et al., 2010) |
| **LV EDV** | (Berko et al., 1987; Carabello et al., 1989, 1991; Dell'Italia et al., 1995, 1997; Katayama et al., 1988; Kleaveland et al., 1988; J. D. Lee et al., 1985; Liu et al., 2013; Matsuo et al., 1998; Nakano et al., 1990, 1991; Nemoto et al., 2002, 2005; Pat et al., 2008, 2010; Perry et al., 2002; Spinale et al., 1993; J. A. Stewart et al., 2003; Trappanese et al., 2015; Tsutsui et al., 1994; Urabe et al., 1992; Zheng et al., 2009) | -- |
| **Myocyte Cell Area/Length** | (Dell'Italia et al., 1997; Kihara et al., 1988; Pat et al., 2008; Perry et al., 2002; Spinale et al., 1993; Tsutsui et al., 1994; Urabe et al., 1992) | (Du et al., 2010; Jobe et al., 2009) |
| **Concentration in plasma of hypertrophy-related hormones and enzymes** | | |
| **ANP** | (Asano et al., 1999, 2001; Hori et al., 2010; Ichiki et al., 2013) | (Abassi et al., 2011; Huang et al., 1992; Langenickel et al., 2000; Pagel et al., 2002; Willenbrock et al., 1997) |
| **BNP** | (Asano et al., 1999, 2001; Hori et al., 2010; Ichiki et al., 2013) | (Langenickel et al., 2000; Wang et al., 2012) |
| **ANGII** | (Ichiki et al., 2013; Perry et al., 2002; Tallaj et al., 2003) | (Bauer et al., 1997; Ishiye et al., 1995; Pagel et al., 2002; Shuji et al., 2000; Suzuki et al., 2004; Wang et al., 2012) |
| **NE** | (Hankes et al., 2006; J. D. Lee et al., 1985; Nagatsu et al., 1994; Tsutsui et al., 1994) | (Huang et al., 1992; Kristen et al., 2002, 2006; Oka et al., 1993; Suzuki et al., 2004; Willenbrock et al., 1997) |
| **ET1** | (Cavero et al., 1990; Ray et al., 2008) | (Gardner et al., 2010; Ishiye et al., 1995) |
| **Intracellular signaling protein activity/phosphorylation** | | |
| **FAK** | (Pat et al., 2010; Sabri et al., 2008) | (Seqqat et al., 2012) |
| **Akt** | (Sabri et al., 2008) | (Dent et al., 2007) |
| **ERK12** | (Sabri et al., 2008) | (Kolpakov et al., 2009; Zhang et al., 2010) |
| **ERK5** | (Liu et al., 2013) | -- |
| **JNK** | (Sabri et al., 2008) | (Kolpakov et al., 2009) |
| **p38** | (Liu et al., 2013; Sabri et al., 2008) | (Kolpakov et al., 2009) |
| **STAT** | -- | (Kolpakov et al., 2009) |
| **cGMP** | (Liu et al., 2013; Trappanese et al., 2015) | (Arnal et al., 1993; Pagel et al., 2002) |
| **ELK** | -- | (Zhang et al., 2010) |
| **Protein abundance** | | |
| **ANP** | | (Fareh et al., 1996; Lachance et al., 2014; Langenickel et al., 2000; Yamakawa et al., 2000) |
| **BNP** | (Zheng et al., 2009) | (Fareh et al., 1996; Langenickel et al., 2000; Yamakawa et al., 2000) |
| **αMHC** | (Imamura et al., 1994; Matsuo et al., 1998) | (Freire et al., 2007; Lachance et al., 2014; Pu et al., 2009). |
| **βMHC** | | |
| **SERCA2** | (Zheng et al., 2009) | (Pu et al., 2013; Wojciechowski et al., 2010) |

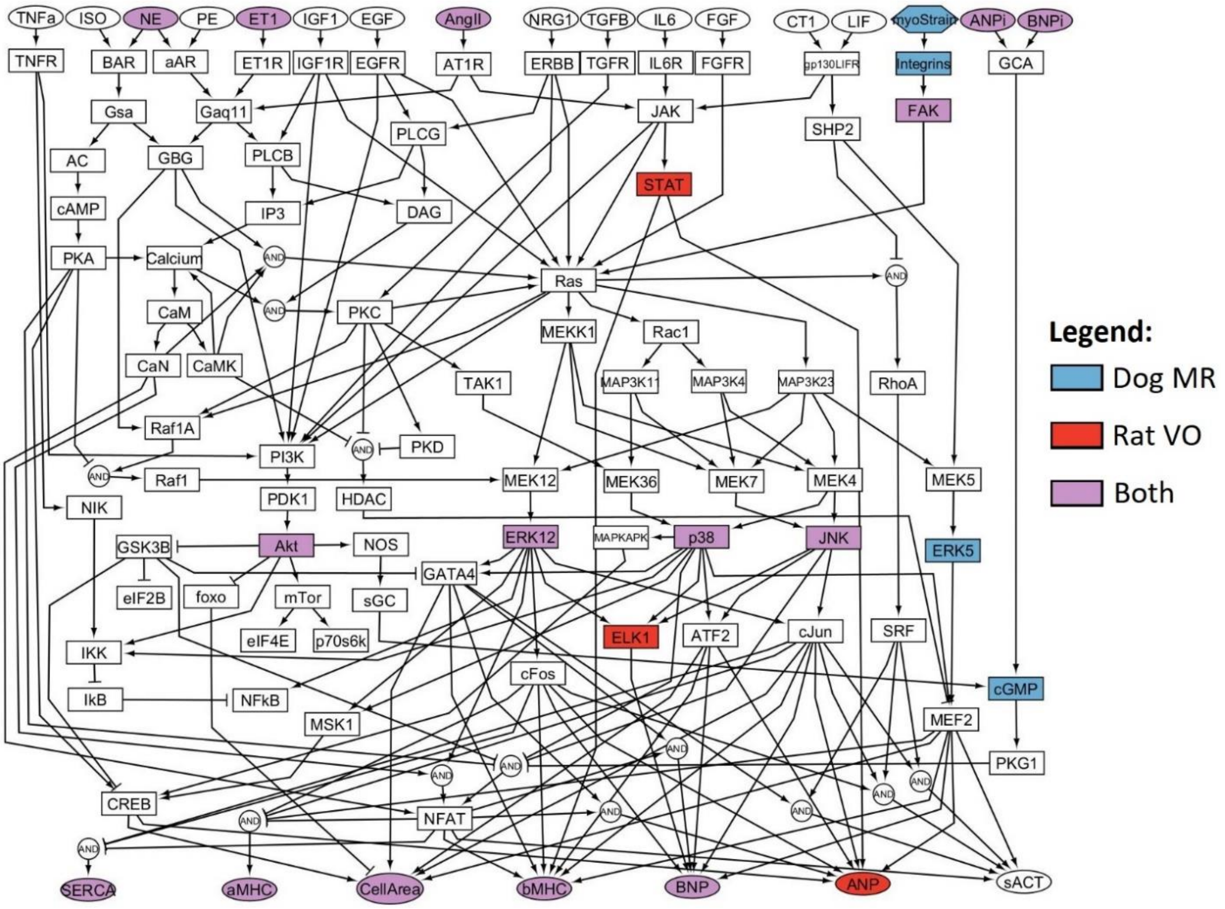

**Figure S1.1.** Location in the network signaling model of available experimental data. See specific references in table S1.1.

**Table S1.2.** Data sources referenced in validation and discussion, not used for calibration.

| Stimulus/Drug | Animal Model | Source |
|---|---|---|
| **NE** | Dog | (King et al., 1987; Laks et al., 1973; Raum et al., 1983; J. M. Stewart et al., 1992) |
| **ANGII** | Rat | (Baltatu et al., 2000; Cassis et al., 1998; Dilley, 1998; Dostal & Baker, 1992; Fabris et al., 2007; Goldspink et al., 2001; Griffin et al., 1991; Grobe et al., 2007; Herizi et al., 1998; Kim et al., 1995; Mishra et al., 2019; Van Eekelen & Phillips, 1988; Wu et al., 2004; Zou et al., 1996) |
| **ISO** | Rat | (Al-Rasheed et al., 2015; Boluyt et al., 1995; Brand et al., 1993; Chowdhury et al., 2013; Ennis et al., 2003; Golomb et al., 1994; Kitagawa et al., 2004; Leenen et al., 2001; Miyoshi et al., 2019; Murad & Tucci, 2000; Nagano et al., 1992; Takeshita et al., 2008) |
| **ISO+βB** | Rat | (Boluyt et al., 1995; Grimm et al., 1999; Hanada et al., 2008; Kitagawa et al., 2004) |
| **ISO+ARB** | Rat | (Golomb et al., 1994; Goyal & Mehta, 2012; Grimm et al., 1999; Leenen et al., 2001; Miyoshi et al., 2019) |
| **NE+ERA** | Rat | (Kaddoura et al., 1996) |
| **AngII+ERA** | Rat | (Ficai et al., 2001; Herizi et al., 1998; Park & Schiffrin, 2002) |
| **VO+βB** | Dog | (Pat et al., 2008; Sabri et al., 2008; Trappanese et al., 2015) |
| | Rat | (Kobayashi et al., 2008; Seqqat et al., 2012) |
| **VO+ARB** | Dog | (Perry et al., 2002) |
| | Rat | (Ishiye et al., 1995; Ruzicka et al., 1993; Zhang et al., 2010) |
| **VO+ERA** | Rat | (Francis et al., 2004; D. S. Lee et al., 2005; Murray et al., 2008, 2009) |
| **VO+ERA+ARB** | Rat | (Fareh et al., 1996; Leskinen et al., 1997) |

### S1.2. Time-varying curves of neurohormonal alterations

We fitted the time course of variations in serum concentrations by allowing a step change immediately after the onset of VO and then fitting a continuous function over the remainder of the time course. Visual inspection of integrated experimental data suggested that linear functions were sufficient to fit the time-varying fold changes of NE, ANP, and BNP beyond an initial step (if present). Beyond an initial step increase, fold changes in AngII concentration appeared consistent with an exponential decay. Finally, no discernable trend was observed in ET1 circulating concentration beyond the initial step, so we fitted for a single constant level for the remainder of the time course. We estimated the probability distribution of each function parameter with independent MCMC runs for each hormone, using the reported experimental concentrations to estimate the likelihood of each set of parameters.

**Table S1.3.** PDFs for fitted parameters of functions describing neurohormonal alterations in VO.

| Species | **Function** | **a** | **b** | **C** |
|---|---|---|---|---|
| ANP | $\frac{w_{ANP}}{w_{ANP}{}^0} = at/\tau + b$ | $1.0747 \pm 0.5612$ | $3.8311 \pm 2.0542$ | -- |
| BNP | $\frac{w_{BNP}}{w_{BNP}{}^0} = at/\tau + b$ | $1.5746 \pm 0.2508$ | $1.05 \pm 0.05$ | -- |
| ET1 | $\frac{w_{ET1}}{w_{ET1}{}^0} = a$ | $3.0 \pm 0.98$ | -- | -- |
| NE | $\frac{w_{NE}}{w_{NE}{}^0} = at/\tau + b$ | $0.1213 \pm 0.0802$ | $1.4610 \pm 0.211$ | -- |
| AngII | $\frac{w_{AngII}}{w_{AngII}{}^0} = ae^{-t/b} + c$ | $4.46 \pm 0.98$ | $0.533 \pm 0.178$ | $2.757 \pm 0.793$ |

### S1.3. Estimation of time-varying probability distributions of stretch

In a spherical model of the left ventricle, end-diastolic stretch can be calculated from the unloaded volume $V_0$ and the end-diastolic volume $V_{ED}$ (see Section 2.4). If the LV is growing and remodeling, both of these volumes change over time. $V_{ED}$ is typically measured and reported in overload experiments, while $V_0$ must be estimated from $V_{ED}$ and other information. Designating individual time steps using the superscript *i*, this section outlines how we used available data to generate probability distributions for the time-varying values of end-diastolic and unloaded ventricular volumes $\left(V_{ED}^i, V_0^i\right)$ in dogs experiencing volume overload.

Step 1: We estimated the PDFs of unloaded ventricular dimensions at baseline, prior to the onset of volume overload. For this, we used reported data on end-diastolic volume, thickness, and maximum fiber stretch $(V_{ED}^0, h_{ED}^0, \lambda_f^0)$ to calculate the unloaded volume and thickness at baseline $(V_0^0, h_0^0)$ using the definition of stretch (Equation 2, main manuscript), the geometry of a thin-walled sphere, and the assumption that the ventricular wall is incompressible, with LVM being written at any time step i (including i=0) as

$$LVM^i = \rho\left[\left(r_{ED}^i + h_{ED}^i\right)^3 - \left(r_{ED}^i\right)^3\right] = \rho\left[\left(r_0^i + h_0^i\right)^3 - \left(r_0^i\right)^3\right] \qquad \text{(Equation S1.1)}$$

where $\rho$ is the density of the myocardium that we assume to be constant. We used unrestricted random sampling of the inputs $(V_{ED}^0, h_{ED}^0, \lambda_f^0)$ to estimate the PDFs of the outputs $(V_0^0, h_0^0)$ over 100,000 iterations using the conventional Monte Carlo method (Figure S1.2).

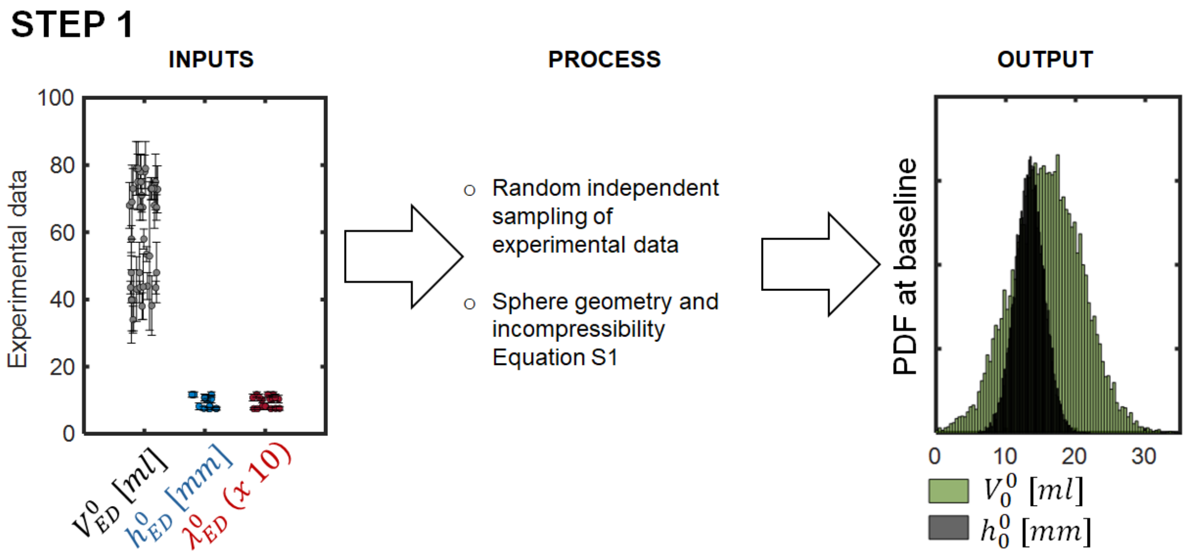

**Figure S1.2.** Estimation of PDF of baseline unloaded ventricular dimensions from available experimental data from baseline end-diastolic dimensions.

Step 2: We estimated time-varying continuous PDFs of $LVM$ and $V_{ED}$ over the course of VO. For this, we assumed that changes in LV mass and end-diastolic diameter follow an exponential rise, consistent with the data shown in Figure 3:

$$\frac{LVM^i}{LVM^0} = C_M\left(1 - e^{\frac{t^i}{\tau_M}}\right) + 1 \qquad \text{Equation S1.2}$$

$$\frac{V_{ED}^i}{V_{ED}^0} = \begin{cases} 1 & at\ \ t^i = 0 \\ C_V\left(1 - e^{\frac{t^i}{\tau_V}}\right) + D_V & at\ \ t^i > 0 \end{cases} \qquad \text{Equation S1.3}$$

where $t^i$ is the time at growth step i, and $C_M$, $C_V$, $\tau_M$, $\tau_V$, and $D_V$ are empirical parameters fitted to $\frac{LVM^i}{LVM^0}$ and $\frac{V_{ED}^i}{V_{ED}^0}$ data derived from experimental measurements. Across multiple studies, larger animals with larger hearts at baseline will also have larger hearts following VO. We calculated the correlation between starting and ending masses and volumes during VO (PCC=0.87) using individual dog data reported by Ross et al. (1972) and Badke and Covell (1979). We then used the method of Hayya et al. (1975) for the division of two correlated and normally distributed variables to derive data on fold changes with respect to baseline $\left(\frac{LVM^i}{LVM^0}, \frac{V_{ED}^i}{V_{ED}^0}\right)$ from available experimental data $\left(LVM^0, V_{ED}^0, LVM^i, V_{ED}^i\right)$. We used random unrestricted sampling of exponential model parameters $C_M$, $C_V$, $\tau_M$, $\tau_V$, and $D_V$ to create time-varying curves of fold changes of LVM and $V_{ED}$. On each iteration, these curves (and the associated parameters) were either retained or dropped based on their likelihood with respect to experimental data using the Metropolis-Hasting selection criteria in a classical MCMC setup. The result after 100,000 iterations is the PDFs of parameters $C_M$, $C_V$, $\tau_M$, $\tau_V$, and $D_V$ that most likely reproduce the $\frac{LVM^i}{LVM^0}$ and $\frac{V_{ED}^i}{V_{ED}^0}$ data (Figure S1.3).

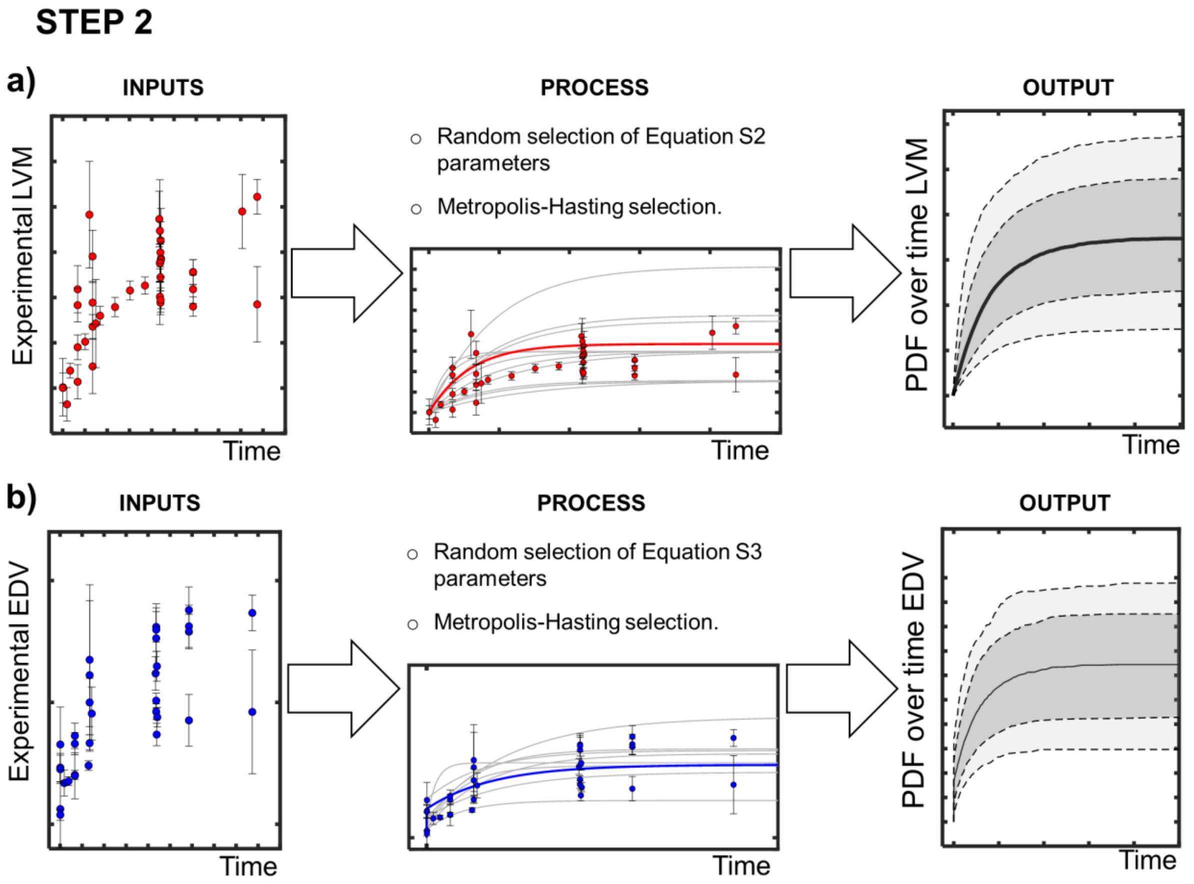

**Figure S1.3.** Estimation of time-varying PDFs for parameters and associated exponential curves describing fold changes of LVM and EDV based on experimental data.

Step 3: Finally, we combined information from the previous steps to estimate the continuous time-varying PDF of stretch over the course of VO. First, we randomly selected a set of exponential parameters $C_M$, $C_V$, $\tau_M$, $\tau_V$, and $D_V$ from their respective PDFs derived in Step 2. Those parameters were either retained or dropped based on the joint likelihood of the asociated chronic $\frac{LVM^i}{LVM^0}$ and $\frac{V_{ED}^i}{V_{ED}^0}$ with respect to experimental data using the Metropolis-Hasting selection criteria. The probability map of $\frac{LVM^i}{LVM^0}$ - $\frac{V_{ED}^i}{V_{ED}^0}$ combinations used to assess that joint likelihood was constructed from 24 experimental datasets from 16 studies reporting both quantities at in experimental MR in dogs (Carabello et al., 1989; Dell'Italia et al., 1995, 1997; Katayama et al., 1988; Kleaveland et al., 1988; Matsuo et al., 1998; Nakano et al., 1991; Nemoto et al., 2005; Pat et al., 2008, 2010; Perry et al., 2002; Spinale et al., 1993; J. M. Stewart et al., 1992; Tsutsui et al., 1994; Urabe et al., 1992; Zheng et al., 2009). As expected, experimentally measured increases in ventricular mass were directly correlated with simultaneously measured increases in end-diastolic volume (PCC=0.56), since both reflect the amount of eccentric hypertrophy that occurred. Following selection of a pair of mass and volume curves, we randomly selected baseline unloaded dimensions $(V_0^0, h_0^0)$ from their PDFs derived in Step 1. We combined the baseline unloaded dimensions and the selected $\frac{LVM^i}{LVM^0}$ time curve to produce a time-varying curve of unloaded volume $\left(V_0^{\mathrm{i}}\right)$ over the course of VO by solving the following equation for $r_0^{\mathrm{i}}$,

$$\frac{LVM^i}{LVM^0} = \frac{\rho\left[\left(r_0^i + h_0^0\right)^3 - \left(r_0^i\right)^3\right]}{\rho[(r_0^0 + h_0^0)^3 - (r_0^0)^3]}, \qquad \text{Equation S1.4}$$

and calculating the unloaded volume $\left(V_0^i = \frac{4}{3}\pi r_0^i\right)$. Equation S1.4 arises directly from the geometry of a sphere (Equation S1.1) with the further assumption that during eccentric hypertrophy, all growth occurs in the radius / circumference of the sphere, with no change in wall thickness. We computed the time-varying curve for stretch for each iteration from the selected $\frac{V_{ED}^i}{V_{ED}^0}$ and computed $V_0^{\mathrm{i}}$ curves using Equation 2 (main manuscript), and generate the time-varying PDF for LV stretch by repeating this entire process over 100,000 iterations (Figure S1.4).

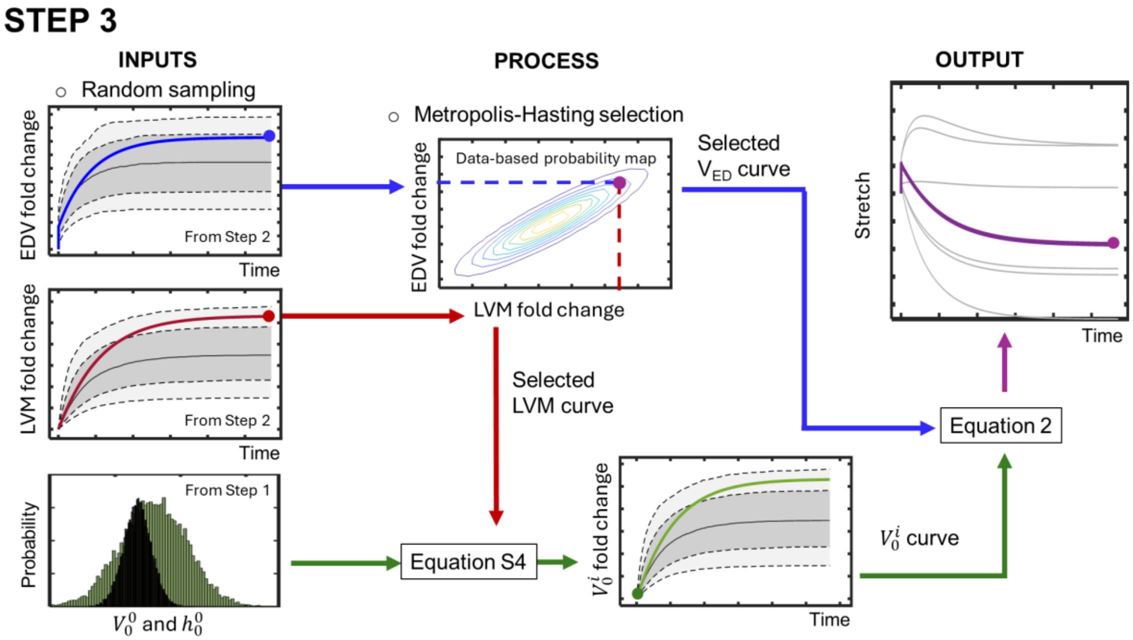

**Figure S1.4.** Estimation of the time-varying probability distribution of ventricular stretch from PDFs of LVM, EDV and baseline unloaded dimensions

The experimental correlations between baseline and chronic dimensions (employed in Step 2) (Badke et al., 1979; Ross & McCullagh, 1972) and between EDV and LVM growth (employed in Step 3) (Carabello et al., 1989; Dell'Italia et al., 1995, 1997; Katayama et al., 1988; Kleaveland et al., 1988; Matsuo et al., 1998; Nakano et al., 1991; Nemoto et al., 2005; Pat et al., 2008, 2010; Perry et al., 2002; Spinale et al., 1993; J. M. Stewart et al., 1992; Tsutsui et al., 1994; Urabe et al., 1992; Zheng et al., 2009) are shown graphically in figure S1.5.

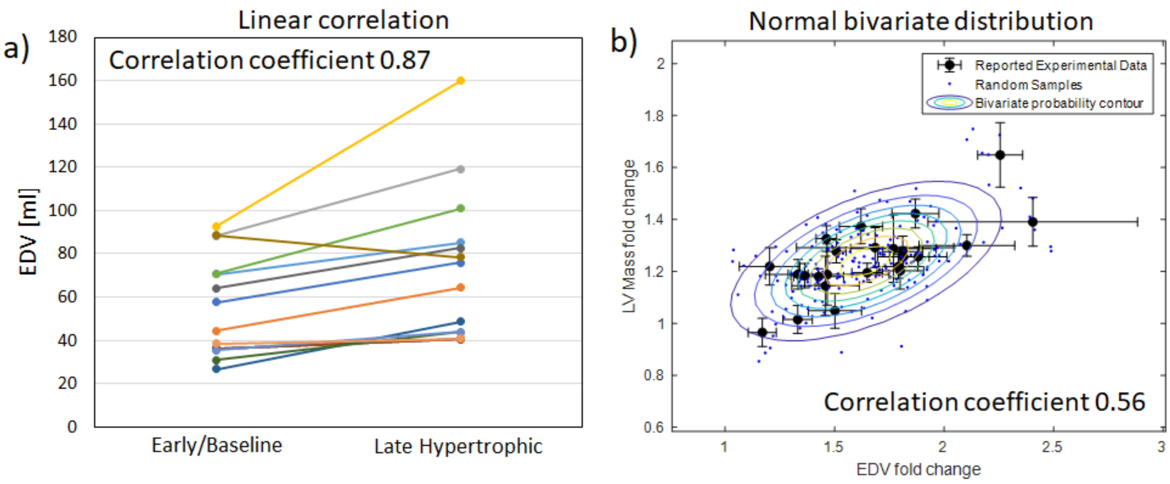

**Figure S1.5.** Correlations considered in the Bayesian analysis of changes in LV stretch during VO. a) Correlation between baseline and chronic- volumes during experimental VO in dogs. b) Co-variance of growth as assessed by changes in ventricular mass and cavity volume from coupled data of experimental MR in dogs.

#### S1.4. Implementation of the Markov Chain Monte Carlo algorithm

The Markov Chain Monte Carlo (MCMC) algorithm with Metropolis Hasting selection criteria is a standard tool for Bayesian statistical analysis and machine learning. The algorithm utilizes known evidence of the behavior of a system to produce probability distributions for the model parameters (Figure S1.6). The quality of the outputs of MCMC depends on the number of iterations of the chain. The number of iterations required to reach convergence depends on the specific problem. To obtain reliable and reproduceable outputs we implemented the algorithm in two stages. In the first stage we assume a uniform probability distribution for all parameters and run 10,000 iterations. We use the outputs of the first stage as the prior probability distribution for the second stage and perform another 20,000 iterations with checks for convergence every 5,000 iterations. Convergence was reached for all runs before the end of the two-stage process (Figure S1.7).

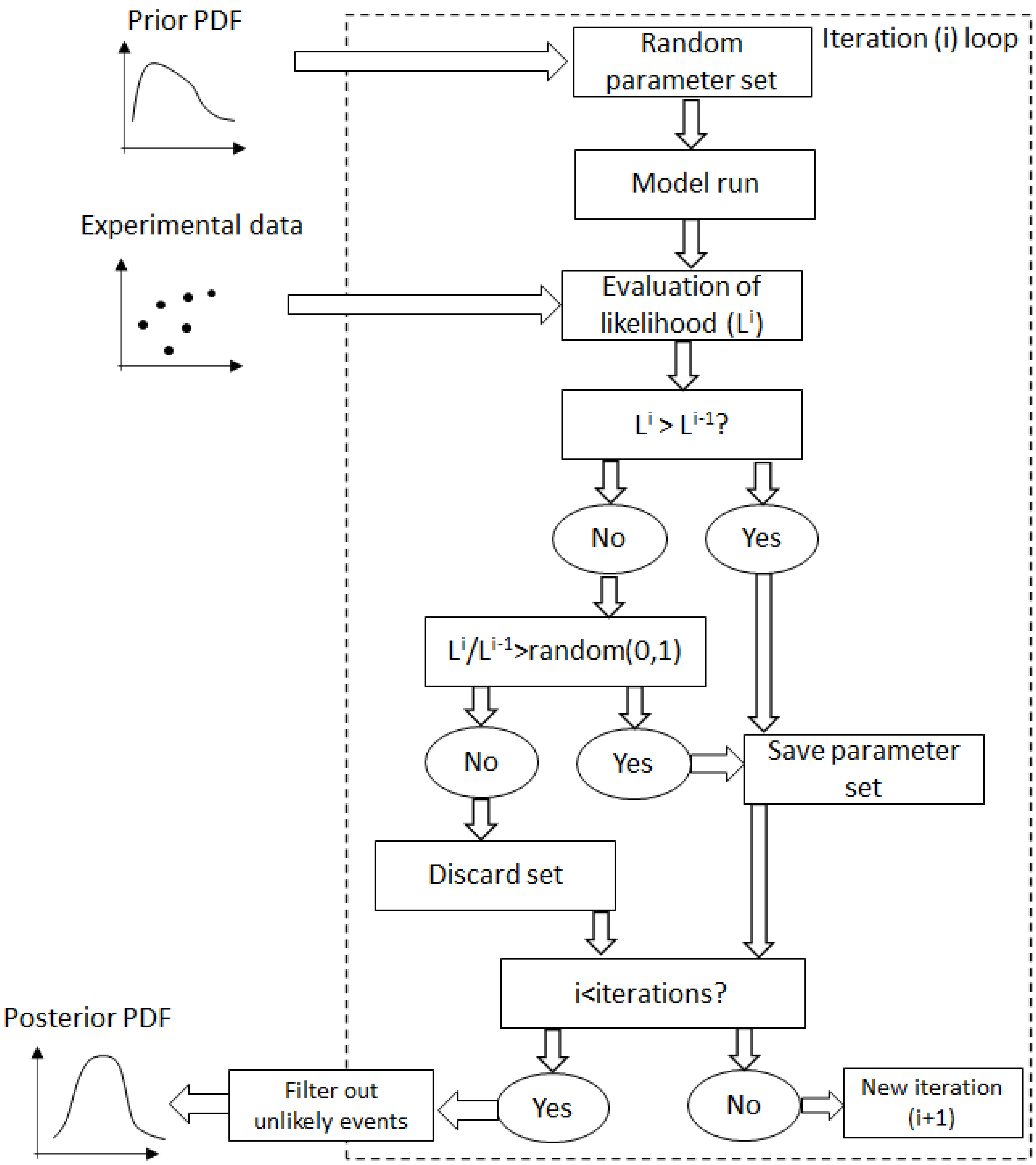

**Figure S1.6.** Diagram of MCM algorithm with Metropolis-Hasting selection criteria.

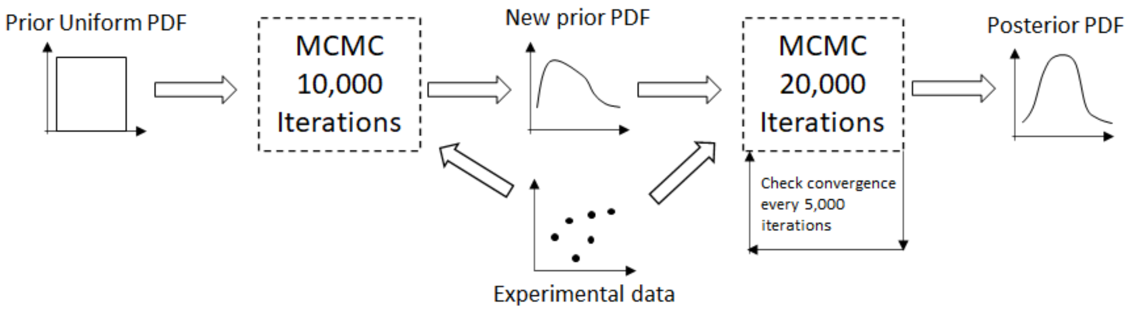

**Figure S1.7.** MCMC implementation in two-stages for reproduceable convergence.

**S1.5. MCMC sampling and posterior parameter probability distributions of network parameters.**

The CellArea node is sensitive to the weight of the myoStrain input, and this parameter heavily influenced the fraction of simulations that produced unlikely scenarios such as reverse growth and runaway growth. By repeating the MCMC analysis for several baseline myoStrain weights $(w^0_{myoStrain})$, we found that too many simulations showed runaway growth driven by ever-increasing NE levels for $w^0_{myoStrain} < 0.05$, while too many solutions showed unphysiologic reversal of growth as strain declined at later time points for $w^0_{myoStrain} > 0.06$. We concluded that $0.05 < w^0_{myoStrain} < 0.06$ is the most likely range for this parameter (Figure S1.8).

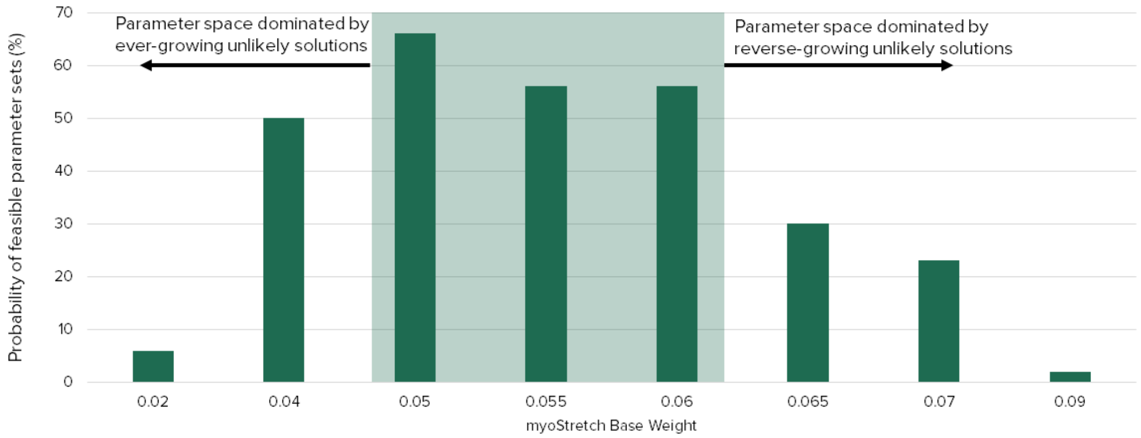

**Figure S1.8.** Percentage of physiologically plausible solutions as a function of myoStrain input weights.

Within the subregion of most likely baseline input weights, each parameter was mostly independent of the others, and displayed a nearly normal probability distribution. The strongest correlation (PCC=-0.42) identified was between the Background and ET1 baseline weights (Figure S1.9).

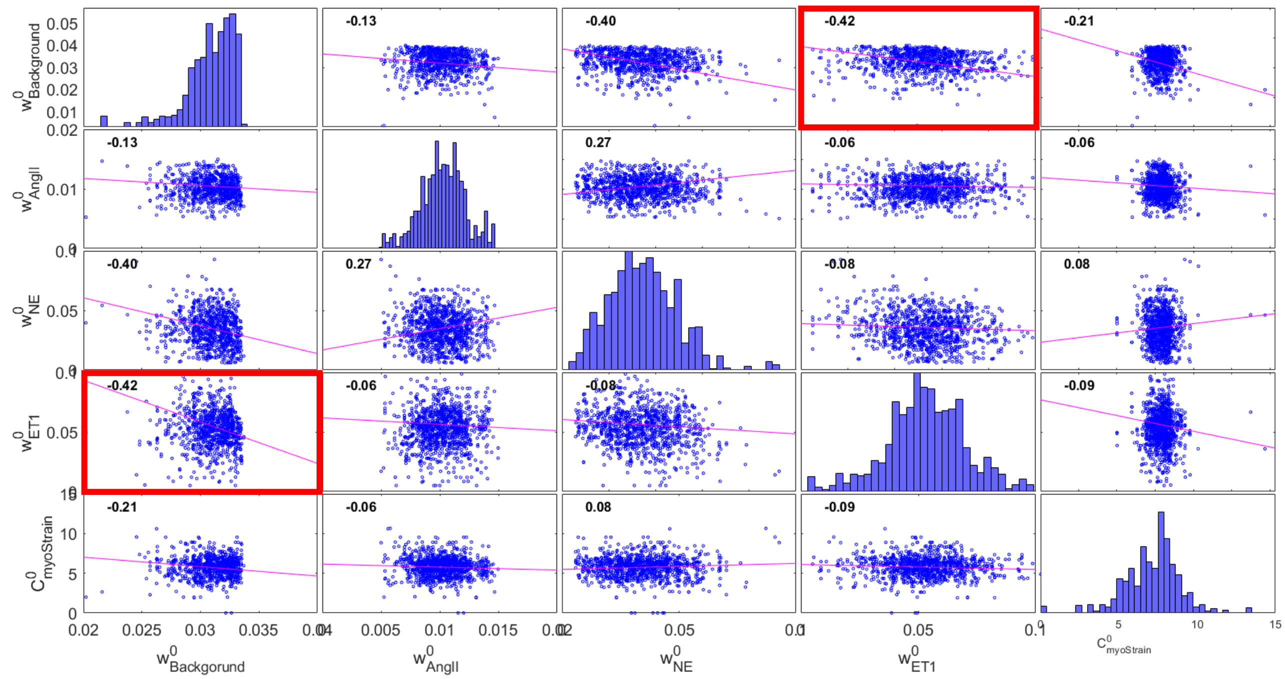

**Figure S1.9.** Correlation matrix of baseline input weights and Strain mapping parameter after filtering-out unlikely results. The Pearson correlation coefficient PCC is shown for each pair of parameters to indicate correlation strength.
